# GREATER EMPATHIC ABILITIES AND THEIR CORRELATION WITH RESTING STATE BRAIN CONNECTIVITY IN PSYCHOTHERAPISTS COMPARED TO NON-PSYCHOTHERAPISTS

**DOI:** 10.1101/2020.07.01.182998

**Authors:** Victor E. Olalde-Mathieu, Federica Sassi, Azalea Reyes-Aguilar, Roberto E. Mercadillo, Sarael Alcauter, Fernando A. Barrios

## Abstract

Psychotherapists constantly regulate their own perspective and emotions to better understand the “other’s” state. We compared 52 psychotherapists with 92 non-psychotherapists to characterized psychometric constructs like, Fantasy (FS) and Perspective Taking (PT), and the emotion regulation strategy of Expressive Suppression (ES), which hampers the empathic response. Psychotherapists showed greater FS, PT and lower ES scores. In a subsample (36, 18 ea.), we did a functional connectivity (FC) study. Psychotherapists showed greater FC between the left anterior insula and the dorsomedial prefrontal cortex; and less connectivity between rostral anterior cingulate cortex and the orbito prefrontal cortex. Both associations correlated with the PT scores and suggest a cognitive regulatory effect related to the empathic response. Considering, that the psychometric differences between groups were in the cognitive domain and that the FC associations are related to cognitive processes, these results suggest that psychotherapists have a greater cognitive regulation over their empathic response.

## INTRODUCTION

Empathy is an essential part of all human social interactions; hence a proper regulation of our empathic response can facilitate better social relations in our daily lives. This is especially true in health care environments, where the relationship between healthcare professionals and patients, has a crucial role in determining therapy success (Goldsmith et al., 2015; Watson et al., 2014). Empathy can be considered as an ‘umbrella’ term that encompasses all processes that emerge, so an observer could understand the “other’s” state by activating their own neural and mental representations of that state. In this view, empathy is a multicomponent process that encompasses cognitive processes, such as mentalizing or emotional regulation, and affective processes such as empathic concern or emotional detection, to name a few (de Waal & Preston, 2017; Decety, 2011; Tousignant et al., 2017; Zaki & Ochsner, 2012). Taking into account the nature of sub-processes that are involved in the empathic response, it is easy to address empathy as a personalized phenomenon which response can be regulated. Thus, diverse sub-process can show differences within sexes or be sensible to training. For instance, women tend to be more empathic than men in relation with affective sub-process, e.g. empathic concern, measure by psychometric, psychological and fMRI tasks (Chrysikou & Thompson, 2016; Mercadillo et al., 2015; Reyes-Aguilar & Barrios, 2016). These differences are also present in the way women suppress their emotions, which is less than men (Flynn et al., 2010). Furthermore, training or professional practice could alter the way we empathized, e.g., loving-kindness expert mediators tend to show more compassion and empathic concern; whereas cognitive perspective taking training could alter mentalizing abilities towards the “other” (Klimecki et al., 2013; Singer & Engert, 2019; Teding van Berkhout & Malouff, 2016).

In a therapeutic environment a proper regulation of the empathic response strengthens the patient-therapist relationship incrementing therapy success (Goldsmith et al., 2015; Teding van Berkhout & Malouff, 2016). Given the dynamic interaction with their patient, psychotherapists need to constantly regulate their empathic response. One way to do this is through the exertion of cognitive control to regulate their own perspective taking and emotional appreciation (Decety, 2011; Ickes, 2016; Lamm et al., 2007; Norcross, John C & Lambert, Michael J, 2019; Rogers, 1992). Part of this, involves avoiding prejudice or rapid judgements, and the use of expressive suppression as an emotional regulation strategy, which hinders empathy (Gross & John, 2003). The constant regulation of such cognitive processes, involves the recruitment of several brain areas, like the left anterior insula (lAi) and the rostral anterior cingulate cortex (rACC). Both areas have been related to diverse empathy sub-processes and emotional regulation strategies. The lAi plays a key role in interoception, forms part of the empathy core network and correlates with affective and cognitive empathy processes, like perspective taking (Fan et al., 2011; Uddin et al., 2017). It has been related to the use of expressive suppression as an emotional regulation strategy (Giuliani et al., 2011; Goldin et al., 2008). The rACC has been associated to emotional regulation, self-regulation, inhibitory emotional control in the use of expressive suppression, and in affective empathy tasks (de Waal & Preston, 2017; Etkin et al., 2015; Kunz et al., 2011).

One way to characterize brain areas interactions, without associating them to a specific stimulus, is by a resting state functional connectivity (FC) study; which is a good first approach when studying cognitive processes that encompass several sub-processes, as in the case of empathy (Guerra-Carrillo et al., 2014). Research has shown that experienced professionals present differences in brain functional connectivity, *e. g.* musicians, meditators (Palomar-García et al., 2017; Taylor et al., 2013). Although, some studies suggest that different types of socio-affective and cognitive training influence changes in the brain (Cohen et al., 2016; Singer & Engert, 2019). To the extent of the reviewed literature, there are no studies that assess the FC of experience professionals of socio-affective and socio-cognitive skills, such as the case of psychotherapists. A characterization of their empathic abilities and use of emotional regulation strategies, could start to shed light into the abilities present in a population, immerse in an environment that requires a constant and dynamic regulation of the empathic response, aimed to strengthen the therapeutic-relationship and thus therapy success. This characterization, from the therapist perspective, could further our understanding of the role of empathy, which is base for a successful patient-therapist relationship.

We hypothesized that if psychotherapists explicitly regulate their empathic response, this regulation will be shown, in differences in their behavioral questioners, between groups and in the expected sex differences, and in their functional connectivity related to cognitive processes of the empathic response, when compare to non-psychoterapists. To test this hypothesis, we applied two behavioral questioners, the Inter Reactivity Index (IRI) (M. Davis, 1980) and the Emotional Regulation Questionnaire (ERQ)(Gross & John, 2003); in a sample of 52 psychotherapists and 92 non-psychotherapists, where group variables of socioeconomic status, sex, age and years of formal studies were controlled. All participants had over 2 years of practice in their respective fields. To evaluate if there were any differences in their functional connectivity we did a voxelwise ROI analysis using lAi and rACC as seeds, in a sub-sample of 36 (18 per group) more experience participants (> 6 years of practice).

## METHODS

### Participants

A sample of 52 psychotherapists (age: 50.1 ± 9 years; 32 women) and 92 non-psychotherapists (age: 52.3 ± 10 years; 41 women) participated in this study; all of them had over 2 years of professional experience and were professionally active. Group variables of age, sex, socioeconomic status and years of formal studies were controlled. Individuals in the psychotherapists group had completed postgraduate studies of psychotherapy. Individuals in the non-psychotherapists group also had completed postgraduate studies, but their studies were from fields of knowledge unrelated to psychotherapy. All participants that agreed to participate signed an informed consent form. For the resting state fMRI (rsfMRI) study, we took a 36 sub-sample of 18 psychotherapists (age: 54.4 ± 7 years; 9 women) and 18 non-psychotherapists (age: 54.6 ± 7 years; 9 women); all of them were right handed, had over 6 years of professional experience and were professionally active. Group variables of age, sex, socioeconomic status and years of formal studies were controlled. The postgraduate studies from the non-psychotherapists group were from seven different fields of knowledge (according to Mexico’s INEGI categorization, Supplemental Material, Table S1), all the professions were unrelated to psychotherapy. Exclusion criteria included neurological disorders, use of psychopharmaceuticals, alexithymia and depression assessed by interview and psychometric tests (Beck depression inventory, BDII; Toronto Alexithimia Scale, TAS-20) (Bagby et al., 1994; Beck et al., 1961); excessive movement during MRI acquisition was also considered within the criteria; none of the participants were excluded. All participants signed an informed consent. Experimental protocols were approved by the institutional bioethics committee and followed the guidelines of the Declaration of Helsinki.

### Psychometric tests

We used the interpersonal reactivity index (IRI) (M. Davis, 1980; M. H. Davis, 1983) to assess both cognitive and affective empathy. These four subscales are: Fantasy (FS), Perspective Taking (PT), Empathic Concern (EC), and Personal Distress (PD), and refer to two cognitive factors (FS, PT) and two affective ones (EC, PD). The IRI is a 28-item self-report questionnaire which responses are measured on a 5-point Likert-type scale ranging from 4 (describes me very well) to 0 (does not describe me well), each of the four subscales are comprised by 7 items, with subscales scored as the sum of the items.

The emotional regulation questionnaire (ERQ) (Gross & John, 2003) was also applied to the sample. The ERQ is a 10-item self-report questionnaire that measures the tendency of an individual to use reappraisal and expressive suppression as emotion regulation strategies. The responses are measure on a seven-point Likert scale ranging from 7 (strongly agree) to 1 (strongly disagree). The ERQ consists of two non-correlated subscales Expressive Suppression (four items) and Reappraisal (six items), with subscales scored as the mean of the items.

### Data Analysis

The test were scored as the respective authors stated (M. Davis, 1980; Gross & John, 2003). For group and gender comparisons, all data was converted to z-values. Differences between groups and intragroup (gender) were evaluated by a two sample t-test. To evaluate the correlations between variables in each group we employed Pearson correlations. To control false positives due to multiple comparisons, false discovery rate (*FDR*) was used in both the two sample t-test and the Pearson correlations. To assess sex effects between groups a two factor (group and sex) ANOVA was applied, to explore the differences due to sex within and between, groups a *post-hoc* Tukey HSD was done.

### Imaging

Brain images were acquired with a 3T MR scanner (General Electric, Waukesha, WI), using a 32-channel array head coil. Whole brain resting state images were acquired using a gradient recalled echo T2* echo-planar imaging sequence (TR= 2000 ms, TE = 40 ms, voxel size 4 × 4 x 4 mm^3^). Participants were instructed to close their eyes during the acquisition. All the scans in this transversal study had the same image sequence parameters, with the exception of the duration of the acquisition, 13 subjects where scan with a duration of 6min, 180 volumes, while the other 23 were scan for 10min, 300 volumes (this was due to the passage of time along the experiment). High resolution structural T1-weighted images were acquired for anatomical reference. Images were acquired using a 3D spoiled gradient recalled (SPGR) acquisition with a 1× 1× 1 mm^3^ voxel size (TR =8.1 ms, TE= 3.2 ms, flip angle =12.0°).

### Image Analysis

Resting state images analysis was performed using in-house scripts and FMRIB’s Software Libraries (FSL v.4.1.9; Jenkinson et al., 2012; Smith et al., 2004). Preprocessing was done using, slice timing correction, inhomogeneity correction, physiological noise and head motion correction, brain extraction, spatial normalization, and high-band-pass temporal filtering (0.01–0.08 Hz). Subsequently, using rigid body transformation, images were registered to the corresponding structural image, then using non-lineal transformations images were normalized to the Montreal Neurological Institute (MNI) standard space. Estimation of motion parameters was done for each volume within the fMRI dataset, and the root mean squares (rms) of the displacement relative to the precedent volume were obtained (Satterthwaite et al., 2013). Participants were to be removed if they had over 30 volumes that showed more than 0.25 mm of rms, none of the participants were discarded. To minimize physiological noise, five principal components of WM and CSF were regressed out, a method termed aCompCor (Behzadi et al., 2007; Chai et al., 2012).

To obtained resting state functional connectivity (FC), a seed analysis of lAi and rACC was performed. Seeds were chosen based on their association to empathy and emotion regulation (Supplemental Material Table S2), described in previous research (de Waal & Preston, 2017; Etkin et al., 2015; Fan et al., 2011; Giuliani et al., 2011; Kunz et al., 2011; Uddin et al., 2017). Both seeds were selected from a NeuroSynth automated meta-analysis executed during February, 2018 (Yarkoni et al., 2011), using the search terms: Cognitive Emotional and Empathy. To verify that the different scan durations wouldn’t affect the results, we first took the 23 subjects of 10 min of duration and did a pair t-test (Winkler et al., 2014) with a version of the first 6min of their scans, comparing their FC of the seeds of interest; there were no differences in connectivity between the two durations. Given this, we shortened the 23 subject’s scans from 10 min of duration to the first 6 min and did with those all the FC analysis. For the construction of the functional connectivity maps, FC was obtained by the Pearson correlation between the seed fMRI temporal series and the temporal series of the different voxels in the whole brain. Afterwards a Fisher’s z transformation was applied. To compare the functional connectivity between the two groups, we first obtained the average functional connectivity map of the two groups for each seed, this was done using one sample t-test for each group. Consequently, we looked for differences between groups within the common network (average map) using a two sample t-test. For the comparison between groups, even that sex as a variable was controlled (9 women in each group), to control de effect of sex within groups, we used sex as a covariate. The p values associated to the t-tests were estimated based on permutation analysis, and significant clusters were identified using family-wise error (FWE) and a Threshold-Free Cluster Enhancement (TFCE; Smith & Nichols, 2009).

## RESULTS

### Psychometric Tests

In the sample of 52 Psychotherapists and 90 non-psychotherapist, the psychotherapists showed higher scores in the cognitive empathy scales of the IRI: Fantasy (FS) and Perspective Taking (PT), when compared with non-psychotherapists. While in the affective empathy scales, Empathy Concern and Personal Distress, there were no differences. Additionally, psychotherapists showed lower scores in the use of Expressive Suppression (ES) measure by the ERQ, whereas non-psychotherapists showed a greater score in the use of such emotional regulation strategy; there were no differences in the use of Reevaluation as strategy (Table 1, Fig.1).

**Fig.1.**
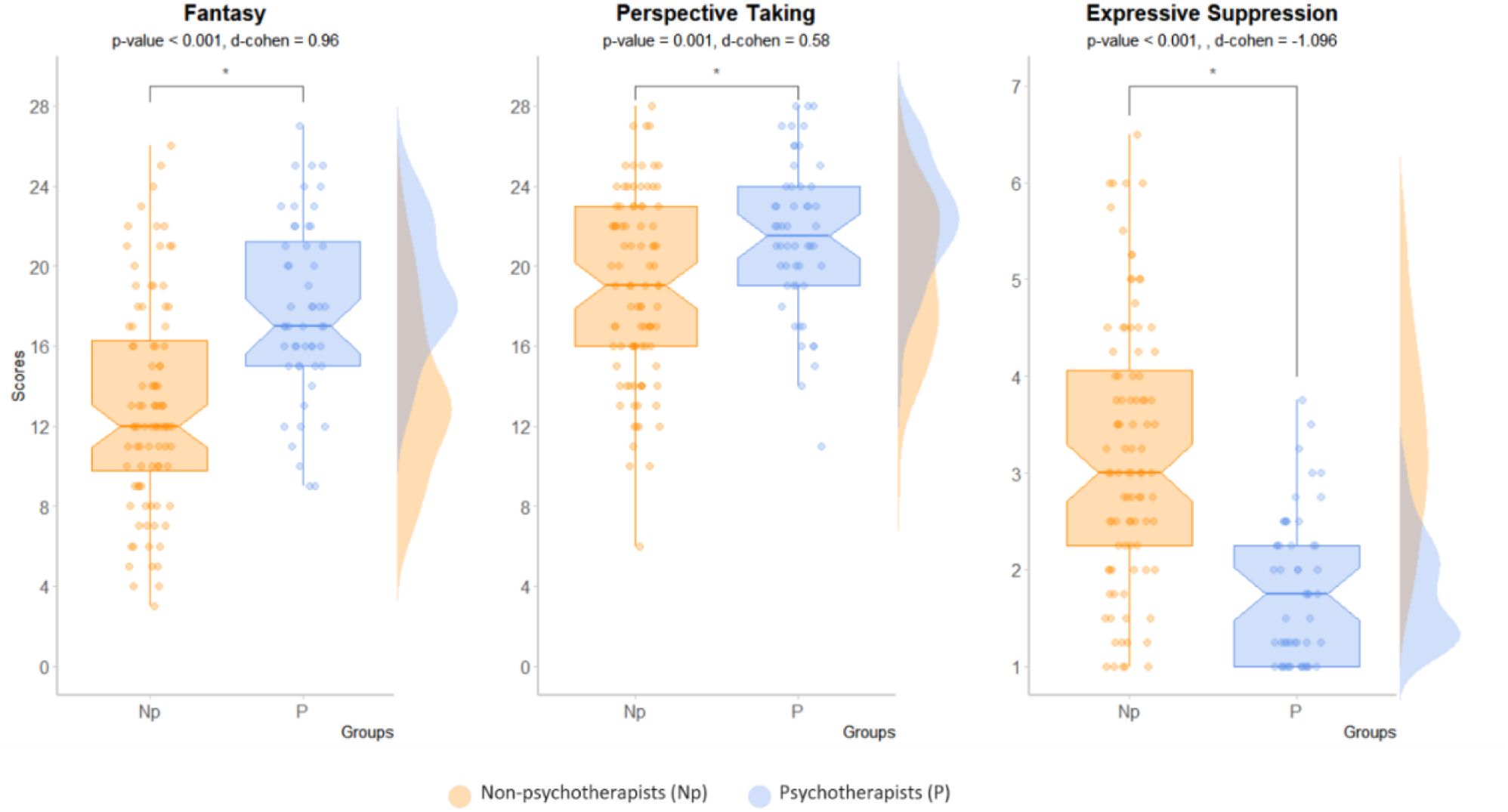
Psychometric differences between psychotherapists and non-psychotherapists. Boxplots of the differences between groups (*FDR-corrected*). In the cognitive empathy scales of the IRI, Fantasy and Perspective Taking, and in the use of Expressive Suppression as a regulation strategy measure by de ERQ. At the right of each boxplot, the density plots of each group are presented. In orange non-psychotherapists (Np), in blue psychotherapists(P). In the y-axis, the scale of the test scores.

**table.1.** Groups Scores.

| Group and Sex | IRI |  |  |  | ERQ |  |
| --- | --- | --- | --- | --- | --- | --- |
|  | FS *<br>(m ± sd) | PT*<br>(m ± sd) | EC<br>(m ± sd) | PD<br>(m ± sd) | RE<br>(m ± sd) | ES*<br>(m ± sd) |
| **Psychotherapists (n=90)** |  |  |  |  |  |  |
| Women | 17.6 ± 5 | 21.5 ± 4 | 21.3 ± 21 | 9.2 ± 4 | 4.4 ± 1 | 1.7 ± 0.8 |
| Men | 18.2 ± 3 | 21.3 ± 3 | 21.0 ± 3 | 8.8 ± 4 | 4.6 ± 1 | 1.8 ± 0.6 |
| Total | 17.9 ± 4 | 21.5 ± 4 | 21.2 ± 3 | 9.0 ± 4 | 4.5 ± 1 | 1.8 ± 0.7 |
| **Non-psychotherapists (n=52)** |  |  |  |  |  |  |
| Women | 13.8 ± 5 | 18.6 ± 4 | 23.3 ± 4 | 10.6 ± 4 | 4.4 ± 1 | 2.8 ± 1 |
| Men | 12.4 ± 5 | 18.9 ± 4 | 20.4 ± 5 | 10.9 ± 5 | 4.5 ± 1 | 3.5 ± 1 |
| Total | 13.0 ± 5 | 18.8 ± 4 | 21.7 ± 4 | 10.8 ± 4 | 4.5 ± 1 | 3.2 ± 1 |

Interpersonal reactivity index (IRI) scales: PT=Perspective Taking, FS=Fantasy, EC=Empathy Concern, PD=Personal Distress. Emotional regulation questionnaire (ERQ) scales: RE=Reevaluation, ES=Expressive Suppression. \*Constructs that showed statistical significant differences (FDR-corrected).

Since the psychotherapists group outperformed in Perspective Taking, Fantasy, and also uses less Expressive Suppression as an emotional regulation strategy, we tested if those differences were also portrayed in the relations between the different test scales, within each group. The psychotherapists group showed a negative correlation between Perspective Taking (PT) and Personal Distress (PD) (Fig.2). Whereas for the non-psychotherapists group, Perspective Taking was positively correlated with Fantasy and Empathic Concern, and these last, Fantasy-Empathic Concern, were also related positively between them. Furthermore, Expressive Suppression which was higher in non-psychotherapist showed a negative correlation with Empathic Concern in this group.

**Fig.2.**
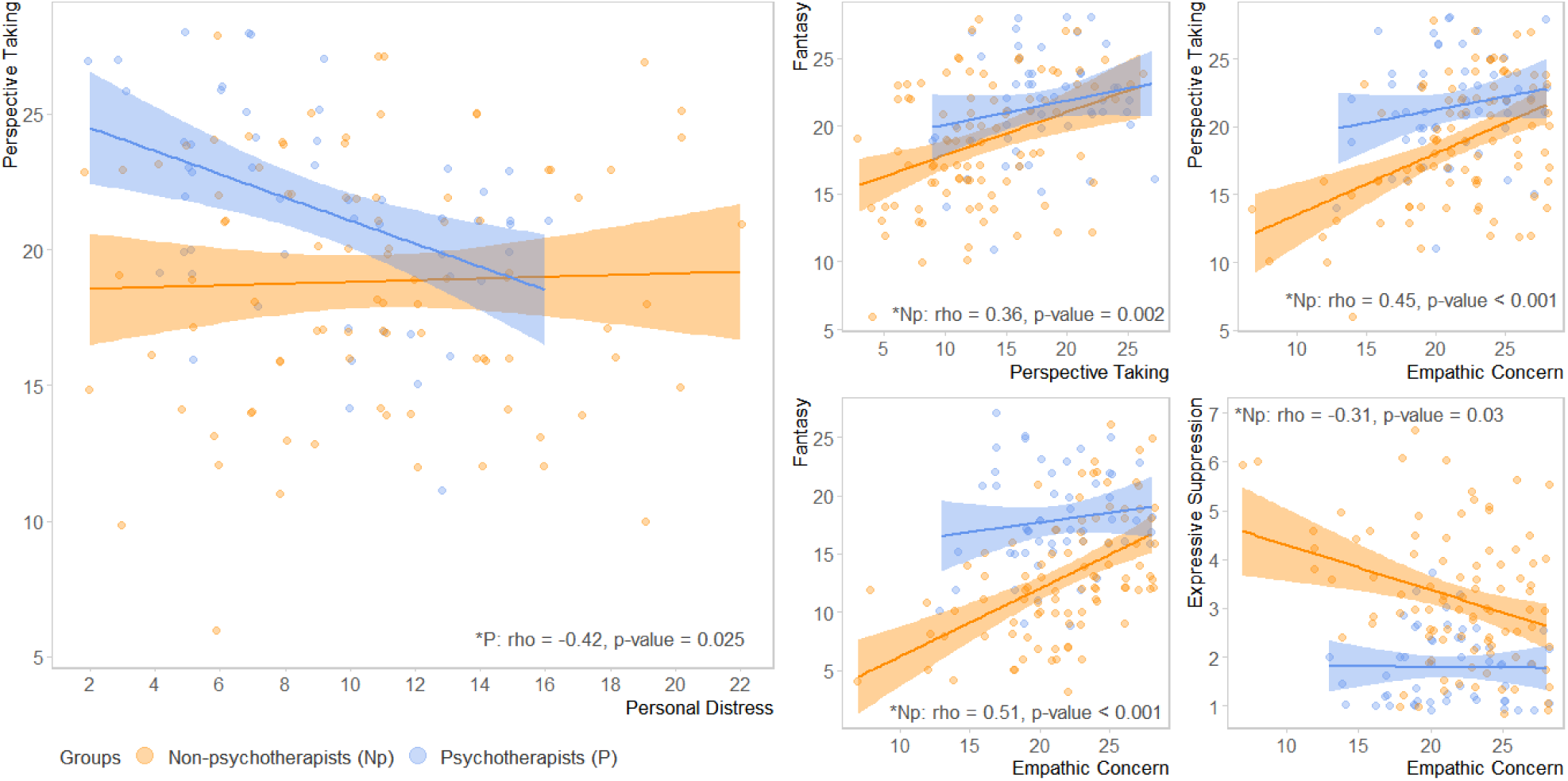
Correlations between constructs within each group. In the graph on the left the significant correlation between PT and PD showed in the psychotherapist group. At the right the four significant correlations showed in the non-psychotherapists group. All p-values are *FDR-corrected*. In orange non-psychotherapists (Np), in blue psychotherapists(P). In the “x” and “y” axis, the scale of the test scores.

#### Sex Differences

There were no sex differences in the psychotherapists group, the sex differences were only present in the non-psychotherapists. These differences were in the scales of Empathy Concern and Expressive Suppression (Table 2). Where woman from the non-psychotherapists group showed higher Empathy Concern and lesser use of Expressive Suppression, in respect to non-psychotherapists men. Any influence of sex in the differences presented between the two groups (NP and P) was discarded by a *post-hoc* analysis (Fig.3).

**Fig.3.**
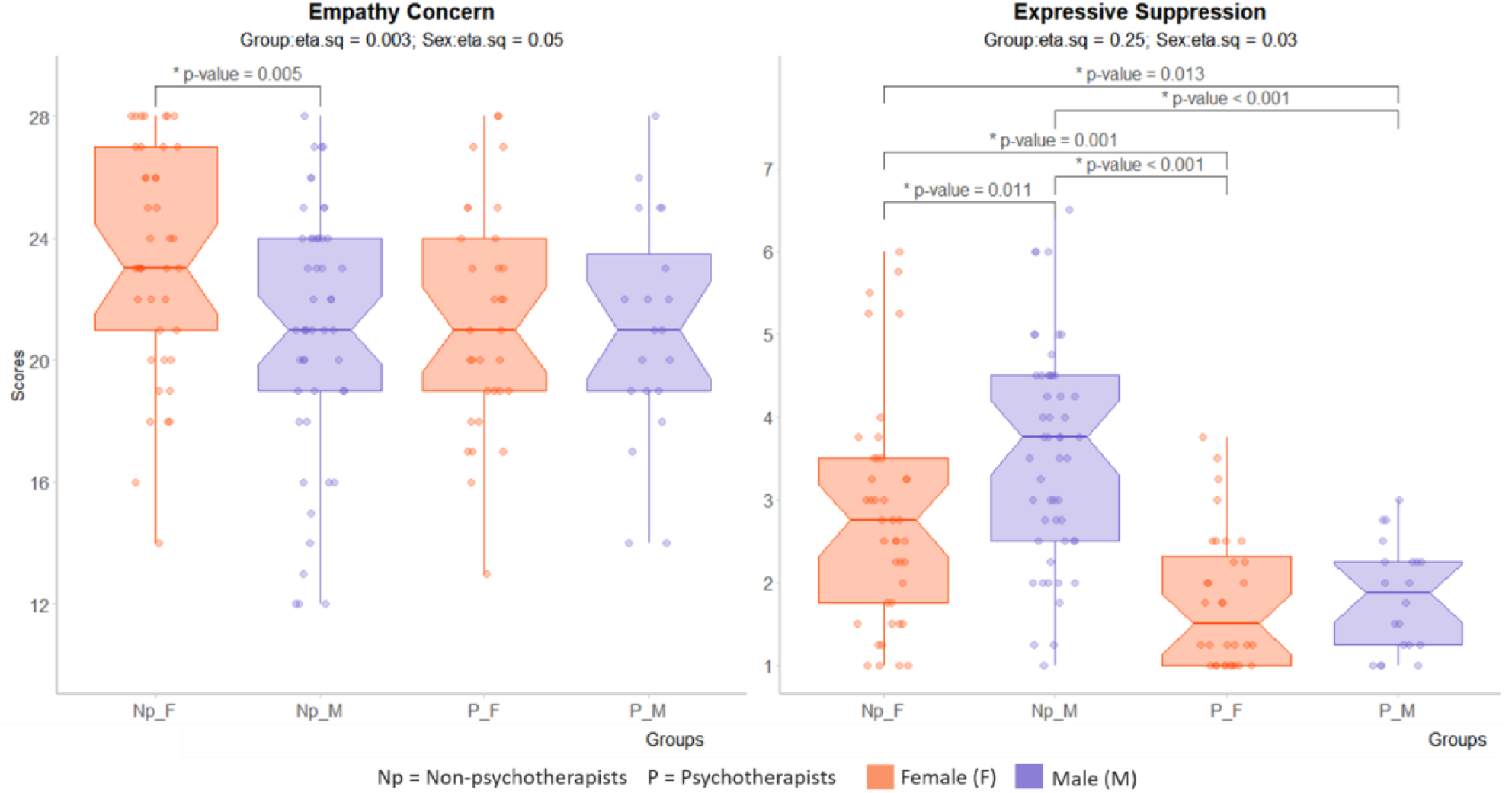
Sex differences *post-hoc* analysis. In the graph of the left the differences between sex in the Empathy Concern Scale and in the graph of the right in the construct of Expressive Suppression. In red females (F), in purple males (M). In the y-axis, the scale of the test scores.

**Table 2.** Group and sex differences, ANOVA results.

| Construct | Group difference | Sex difference | Group-Sex Interaction* |
| --- | --- | --- | --- |
| **Expressive Suppression** | p < 0.001 | p = 0.009 | - |
| **Empathic Concern** | - | p = 0.004 | - |
| **Fantasy** | p < 0.001 | - | - |
| **Perspective Taking** | p < 0.001 | - | - |

### rsfMRI

The seed based FC analysis showed differences when contrasting psychotherapists and non-psychotherapists (Table 3, Fig.4). Psychotherapists showed greater connectivity between the left anterior insula (lAi), associated with the empathy core network, and the dorsomedial prefrontal cortex (dmPFC). Conversely, psychotherapists showed lesser connectivity between the rostral anterior cingulate cortex (rACC), associated with emotional regulation and expressive suppression, and the orbito prefrontal frontal cortex (oPFC); in respect to non-therapists (Fig. 4, Table 3).

**Fig.4.**
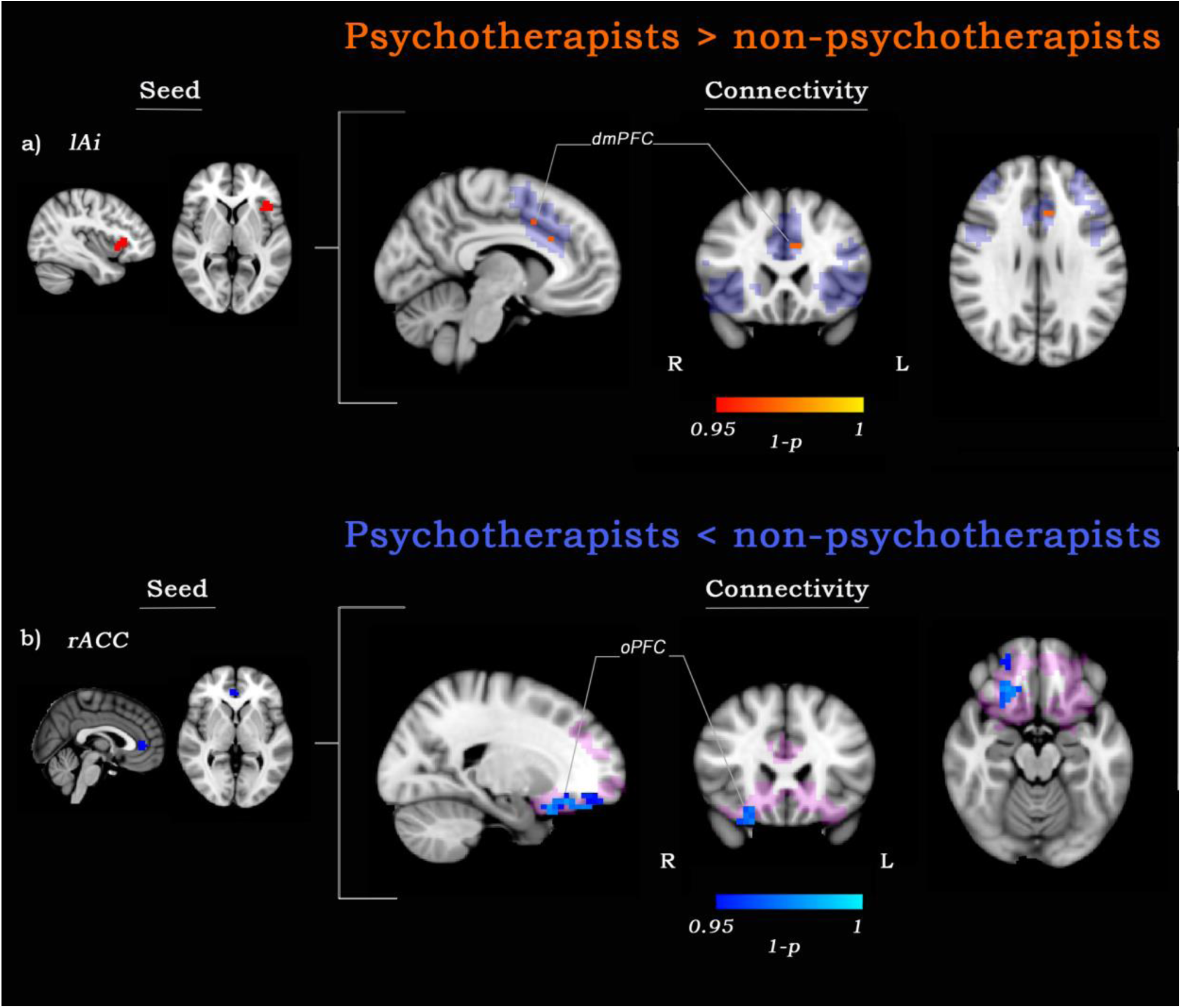
Functional connectivity differences. (a) Seed left anterior insula (lAl) (red), grater connectivity with dorsomedial prefrontal cortex (dmPFC) (red-yellow; 1-p-value > 0.95 *FWE corrected*). In purple, conjunction mask of average FC maps for both groups. (b) rostral anterior cingulate cortex (rACC) (blue), lesser connectivity with orbito prefrontal cortex (oPFC) (blue-lightblue; 1-p-value > 0.95 *FWE corrected*). In pink, conjunction mask of average FC maps for both groups.

**Table 3.** FC differences.

| Seed | FC diff. areas<sup>2</sup> | abbr | Cluster | No. Voxels | 1-p-value<sup>3</sup> | MNI Coordinates<sup>1</sup> |  |  |
| --- | --- | --- | --- | --- | --- | --- | --- | --- |
|  |  |  |  |  |  | x | y | z |
| *lAi* | *Dorsomedial prefrontal cortex* | *dmPFC* | 2 | 2 | 0.97 | -10 | 22 | 28 |
|  |  | *dmPFC* |  | 1 | 0.96 | -6 | 10 | 40 |
| *rACC* | *Orbito prefrontal cortex* | *oPFC* | 3 | 45 | 0.981 | 22 | 18 | -20 |
|  |  | *oPFC* |  | 1 | 0.951 | 14 | 42 | -20 |
|  |  | *sgACC* |  | 1 | 0.952 | 6 | 30 | -8 |

<sup>1</sup>Peak with the maximum p-value, <sup>2</sup>Brain areas that which FC with the seed showed differences, <sup>3</sup>All the p-values are *FEW corrected*.

### FC and cognitive empathy

The psychotherapists of the rsfMRI study showed the same psychometric differences when compare with the non-psychotherapists sub-group, shown in the bigger sample (Supplemental Material, Fig.S1). The psychotherapists, who outperformed in Perspective Taking as a cognitive empathy measure by the IRI, showed a negative correlation between Perspective Taking and their FC of the lAi seed with the dmPFC (lAi-dmPFC), and the rACC seed with the oPFC (rACC-oPFC) (Fig.5).

**Fig.5.**
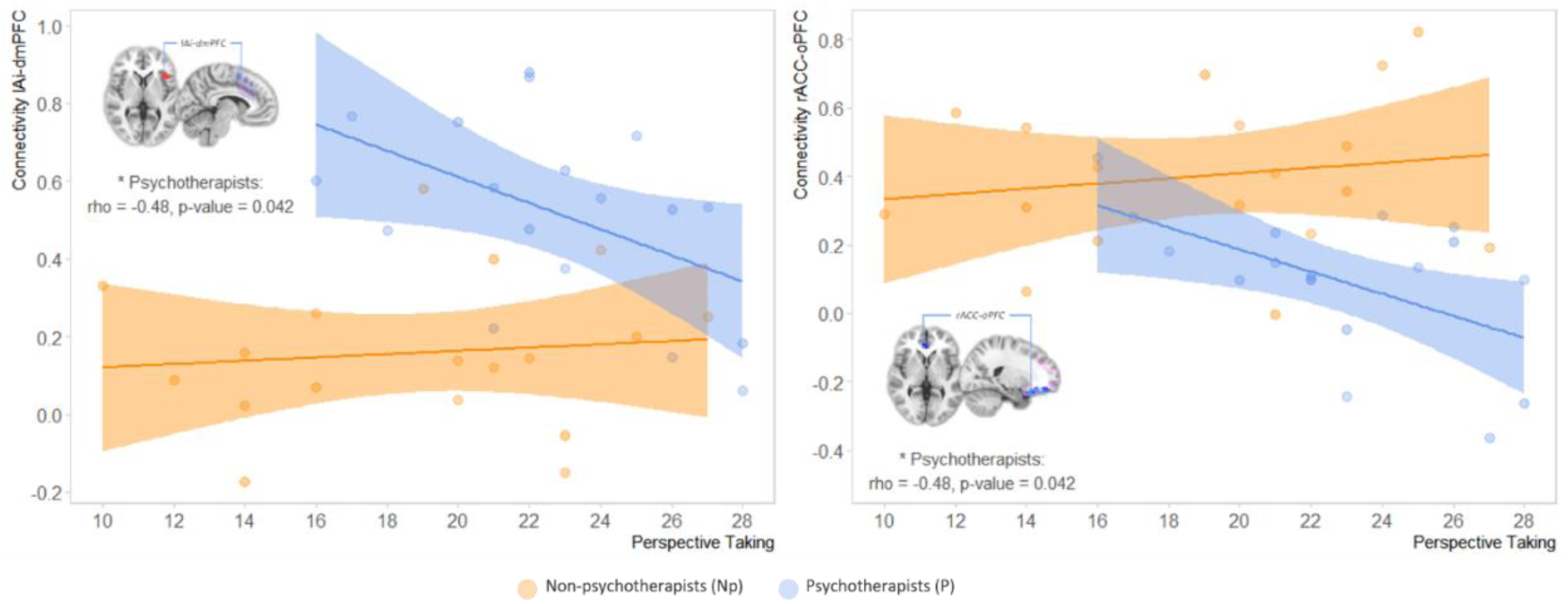
Correlation between Functional connectivity (FC) and Perspective Taking scores. In the graph of the left the correlation between Perspective Taking and the FC of the the left anterior insula with dorsomedial prefrontal cortex (lAi-dmPFC). In the graph of the right the correlation between Perspective Taking and the FC of rACC with orbito prefrontal cortex (rACC-oPFC). p-values are *FDR-corrected*. In orange non-psychotherapists (Np), in blue psychotherapists(P). In the “x”, the scale of the test scores and “y” axis the FC values.

## DISCUSSION

The empathic response, as an umbrella construct, can be regulated by practice or training. In this study, psychotherapists when compared to non-psychotherapists, showed greater scores in Fantasy (FS) and Perspective Taking (PT), both are cognitive-empathy constructs, that refer, in layman’s terms, “to put yourself in the other’s shoes” (Shamay-Tsoory, 2011). These differences might be related to their professional practice, psychotherapists have to constantly modulated their perspective to understand more accurate the “other’s” viewpoint (Ickes, 2016; Lamm et al., 2007; Norcross, John C & Lambert, Michael J, 2019; Rogers, 1992). Psychotherapists also showed less use of Expressive Suppression, which is an emotion regulation strategy that inhibits emotional responding behaviors; this inhibition requires greater consumption of cognitive resources, hindering social performance and generating discomfort to others, as a result, it lessens the empathic response, thus, psychotherapists try to avoid the use of such strategy (Butler et al., 2003; Gross & John, 2003; Norcross, John C & Lambert, Michael J, 2019). It could be inferred that psychotherapists implement other strategies that facilitate more their perspective taking.

The high perspective taking showed by the psychotherapist was related to low personal distress or less personal discomfort in an emotional social setting, which relates to social dysfunction and is negative associated to the other IRI scales. Thus, in the psychotherapists group, perspective taking it’s related to the way they emotionally appreciate their social setting (M. H. Davis, 1983); this relation was only present in psychotherapists. The non-psychotherapists showed construct associations that were expected by previous research (M. H. Davis, 1983; Lockwood et al., 2014); due to their high cognitive-empathy and lower use of expressive suppression, psychotherapists showed only a similar non-significant tendency.

Similarly, the expected sex differences were only present in the non-psychotherapists. As has been reported, women tend to have more Empathic Concern, a construct of affective-empathy, and also tend to express more their emotions (Chrysikou & Thompson, 2016; Flynn et al., 2010). In the use of expressive suppression, men from the psychotherapists group showed lower scores of expressive suppression when compare with the non-psychotherapists, regardless of sex. Given that is natural to men to use expressive suppression as strategy, the fact that men psychotherapists use this strategy less than the women from the non-psychotherapists group, suggests that the differences showed by the psychotherapists (men and women) could be owed to a specialization from their training and practice, which involves a more congruent emotional expression for a more accurate empathic response.

Specialization from training and practice, has been related to changes in FC. In our study psychotherapists showed greater FC between lAi and dmPFC, this could suggest a greater top-down processing. The lAi belongs to the empathy core network, which is always active when we represent the affective state of the “other” (Engen & Singer, 2013), it also has been related to the appreciation and integration of external and internal stimuli to process empathy related states (Uddin et al., 2017). On the other hand, the dmPFC has been associated with executive control, emotion regulation success (Etkin et al., 2015; Kohn et al., 2014; Senholzi & Kubota, 2016) and cognitive empathy (Eres et al., 2015). This association between dmPFC and lAi, suggests a greater regulatory control of the empathic response. We could infer, that in the constant practice of regulating the empathic response dmPFC and lAi interact to exert such regulatory control. lAi-dmPFC connectivity related negatively with perspective taking. This inverse relation could be due to the diverse nature of practice-related effects on FC (Kelly & Garavan, 2005), sometimes, once strengthen the connectivity of certain areas, a refinement of such process will occur, resulting in a diminish FC. Thus, within the psychotherapist group, the lesser the connectivity of lAi-dmPFC the higher their perspective taking.

The lesser FC between rACC and oPFC could support the assumption of a greater top down processing. The rACC has been associated with emotional conflict resolution, specifically with implicit-autonomic regulation which is a more rapid emotion regulation, possibly related to experience-dependent alteration of the value of emotion (Etkin et al., 2015). Similarly, research has shown that the rACC codifies personal traits and stereotypes (Delplanque et al., 2019; Heleven & Overwalle, 2019); this characteristic reinforces the notion that the rACC is involve in imminent emotion resolution; stereotypes serve us to promptly react and resolve emotional conflicts, allowing us to make rapid judgments based on known constructs. In turn, the oPFC has been related with impulsive decision making (Hinvest et al., 2011) and has been associated with implicit emotion regulation and motivational reward towards in-group preference (Mauss et al., 2007; Senholzi & Kubota, 2016). Although implicit emotion regulation serves us to achieve an imminent resolution, it could also implied that our resolution will be embedded with our own prejudice; the inhibition of such type of regulation, could decrease prejudice and increase perspective taking. Thus, lesser interaction between rACC and oPFC could suggest the inhibition of hastily resolving emotional conflict in the therapy environment; this hypothesis, might be supported by the negative relation between the FC of rACC-oPFC and higher perspective taking in psychotherapists.

Given that psychotherapists had training and practice in regulating their empathic response, it is possible to think that this factors are associated to their FC of empathy-regulation related networks, resulting in the differences found in our study. Furthermore, the correlation between the scores of PT and the functional associations, could indicate that these networks participate in the regulation of the psychotherapists own perspective taking to get a more accurate understanding of the “other’s” frame of reference. Although the results suggest that training and practice could be influencing these differences, we cannot assert that such differences were present before the psychotherapists began their training, future research could shed light to these temporality aspects.

## ACKNOWLEDGEMENTS

Víctor Enrique Olalde Mathieu is a doctoral student from “Programa de Doctorado en Ciencias Biomédicas”, Universidad Nacional Autónoma de México (UNAM) and has recived CONACyT fellowship no. 330989 (No.CVU: 619655). *This work was supported* by grants from DGAPA-PAPIIT UNAM grant IN203216 (FAB) and CONACyT *grant CB255462 (FAB)*. We thank Leonor Casanova Rico, Leopoldo González-Santos, Ma. de Lourdes Lara Ayala, Juan J. Ortiz and Erick H. Pasaye for their technical support. M.C. Jeziorski for editing of the manuscript. The authors thankfully acknowledge the imaging resources and support provided by the “Laboratorio Nacional de Imagenología por Resonancia Magnética”, CONACyT network of national laboratories.

## ETHICAL STANDARDS

The authors assert that all procedures comply with the ethical standards of the relevant national and institutional committees on human experimentation and with the Helsinki Declaration of 1975, as revised in 2008. The research protocol was revised and accepted by the Bioethics Committee of the Neurobiology Institute, UNAM.

## AUTHOR CONTRIBUTIONS

Victor E. Olalde-Mathieu Fernando A. Barrios and Sarael Alcauter developed the study concept. All authors contributed to the study design. Testing and data collection were performed by Victor E. Olalde-Mathieu, Fernando A. Barrios, Sarael Alcauter and Federica Sassi. Data analysis and interpretation was performed by: Victor E. Olalde-Mathieu, Roberto E. Mercadillo, Azalea Reyes-Aguilar and Sarael Alcauter. Under the supervision of Fernando Barrios and Sarael Alcauter, Victor E. Olalde-Mathieu drafted the manuscript. Azalea Reyes-Aguilar, Roberto E. Mercadillo and Federica Sassi provided critical revisions. All authors approved the final version of the manuscript for submission.

