## Supplementary material for "GREATER EMPATHIC ABILITIES AND THEIR CORRELATION WITH RESTING STATE BRAIN CONNECTIVITY IN PSYCHOTHERAPISTS COMPARED TO NON-PSYCHOTHERAPISTS": Suplemental Tables

| **Table S1. Non-psychotherapists participants distribution according to the different fields of knowledge of their postgraduate studies** | | |
| --- | --- | --- |
| **Fields of knowledge*** | **Psychometric sample** | **rsfMRI sample** |
| Education | 15 | 2 |
| Humanities | 10 | 1 |
| Social Sciences, Administration and Law | 21 | 3 |
| Natural, Computational and Exact Sciences | 19 | 3 |
| Engineering and Construction | 8 | 3 |
| Agronomy and Veterinary | 10 | 3 |
| Health | 9 | 3 |
| **Total** | **92** | **18** |
| *Mexico’s INEGI categorization (INEGI, 2012) | | |

| **Table S2 Empathy related seeds** | | | | | | | | |
| --- | --- | --- | --- | --- | --- | --- | --- | --- |
|  |  |  |  |  | **MNI Coordinates*** | | |  |
| **Seeds** | **abbr** | **Num. Studies** | **Search Terms** | **MetaAnalisis** | **x** | **y** | **z** | **Studies** |
| Left Anterior Insula | lAi | 137 | Empathy | Neurosynth | -40 | 24 | 0 | (de Waal & Preston, 2017; Decety, 2011; Etkin et al., 2015; Fan et al., 2011; Giuliani et al., 2011) |
| Rostral Anterior Cingular Ctx | rACC | 60 | Cognitive Emotional | Neurosynth | -10 | 38 | 6 | (de Waal & Preston, 2017; Decety, 2011; Etkin et al., 2015; Kunz et al., 2011; Lau & Cikara, 2017) |
| * Peak of maximum p-value. | | | | | | | | |

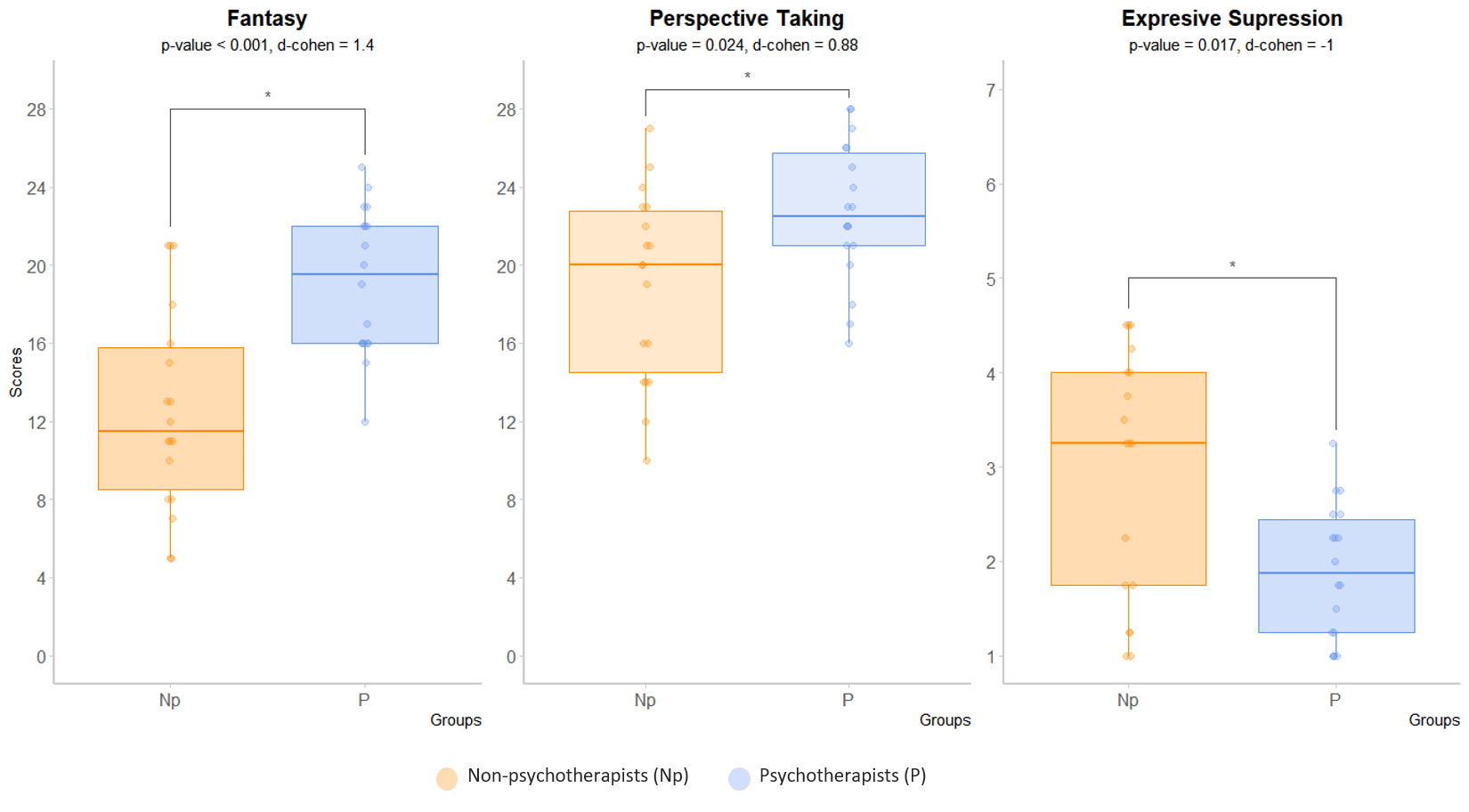

**Fig.S1. rsfMRI sub-sample, psychometric differences between psychotherapists and non-psychotherapists.** Boxplots of the differences between groups (*fdr-corrected*) in the cognitive empathy scales of the IRI, Fantasy and Perspective Taking, and in the use of Expressive Suppression as a regulation strategy measure by de ERQ. In orange non-psychotherapists (Np), in blue psychotherapists(P). In the y-axis, the scale of the test scores.
